# DDGemb: predicting protein stability change upon single- and multi-point variations with embeddings and deep learning

**DOI:** 10.1101/2024.09.05.611455

**Authors:** Castrense Savojardo, Matteo Manfredi, Pier Luigi Martelli, Rita Casadio

## Abstract

The knowledge of protein stability upon residue variation is an important step for functional protein design and for understanding how protein variants can promote disease onset. Computational methods are important to complement experimental approaches and allow a fast screening of large datasets of variations. In this work we present DDGemb, a novel method combining protein language model embeddings and transformer architectures to predict protein ΔΔG upon both single- and multi-point variations. DDGemb has been trained on a high-quality dataset derived from literature and tested on available benchmark datasets of single- and multi-point variations. DDGemb performs at the state of the art in both single- and multi-point variations.

## 1. Introduction

Computational methods for predicting the effect of variations on protein thermodynamic stability play a fundamental role in computational protein design (Notin *et al*., 2024), in functional characterization of protein variants (Vihinen, 2021) and their relation to disease onset (Puglisi, 2022). In the last years, several methods have been presented for the prediction of protein stability change upon variation (ΔΔG).

Tools available can be roughly classified according to the type of information they rely on (protein structure and/or sequence) and on the type of method which carries out the prediction. Structure-based methods rely on the availability of the protein structure in input. Different structure -based predictive approaches have been presented, including methods based on force fields and energy functions (Schymkowitz *et al*., 2005; Kellogg *et al*., 2011; Worth *et al*., 2011), conventional machine-learning methods (Capriotti *et al*., 2005; Dehouck *et al*., 2011; Chen *et al*., 2020; Pires *et al*., 2014b; Laimer *et al*., 2015; Savojardo *et al*., 2016; Montanucci *et al*., 2019), deep-learning approaches (Li *et al*., 2020; Benevenuta *et al*., 2021) and consensus methods (Pires *et al*., 2014a; Rodrigues *et al*., 2018, 2021).

Sequence-based methods only use features that can be extracted from the protein sequence. So far, the vast majority of methods available are based on canonical features such as evolutionary information and physicochemical properties, processed by conventional machine-learning methods (Capriotti *et al*., 2005; Cheng *et al*., 2006; Fariselli *et al*., 2015; Montanucci *et al*., 2019; Li *et al*., 2021). ACDC-NN introduced deep-learning methods (convolutional networks) to process sequence profiles extracted from multiple-sequence alignments (Benevenuta *et al*., 2021). Recently, PROSTATA (Umerenkov *et al*., 2023) adopted protein language models for encoding the protein wild-type and mutated sequences. The protein language model input is then processed in PROSTATA using a simple neural network with a single hidden layer. The sequence based THPLM adopts pretrained protein language models and a simple convolutional neural network (Gong *et al*., 2023).

One of the major challenges in the field of protein stability prediction is the ability to predict ΔΔG upon multi-point variations, i.e. how protein stability is affected when variations occur at multiple residue positions. So far, only a few methods support multi-point variations in input: four structure-based methods (FoldX (Schymkowitz *et al*., 2005), MAESTRO (Laimer *et al*., 2015), DDGun3D (Montanucci *et al*., 2019) and Dynamut2 (Rodrigues *et al*., 2021)), and one sequence-based method, DDGunSeq (Montanucci *et al*., 2019). Overall, the performance of methods for predicting ΔΔG upon multi-point variations is generally lower than that obtained for single-point variations.

In this work, we present a novel method called DDGemb for the prediction of protein ΔΔG upon both single- and multi-point variations. DDGemb exploits the power of ESM2 protein language model (Lin *et al*., 2023) for protein and variant representation in combination with a deep-learning architecture based on a Transformer encoder (Vaswani *et al*., 2017) to predict the ΔΔG.

We train DDGemb using full-length protein sequences and single-point variations from the S2648 dataset (Dehouck *et al*., 2011), previously adopted to train many state-of-the-art approaches (Dehouck *et al*., 2011; Fariselli *et al*., 2015; Savojardo *et al*., 2016).

The performance of DDGemb is evaluated on ΔΔG prediction upon both single- and multi-point variations. For single-point variations, we adopted the S669 dataset recently presented in literature and already adopted for benchmarking a large set of tools (Pancotti *et al*., 2022). For multi-point variations, we adopted a dataset derived from the PTmul dataset (Montanucci *et al*., 2019). In both benchmarks, DDGemb reports state-of-the-art performance, overpassing both sequence- and structure-based methods.

## 2. Materials and Methods

### 2.1. Datasets

#### 2.1.1. The S669 blind test set

For a fair and comprehensive evaluation of DDGemb performance and for comparing with other state-of-the-art approaches, we take advantage of an independent dataset adopted in literature to score a large set of available tools for predicting protein stability change upon variation (Pancotti *et al*., 2022).

The dataset, named S669, comprises 1338 direct and reverse single-site variations occurring in 95 protein chains. ΔΔG values were retrieved from ThermoMutDB (Xavier *et al*., 2021) and manually checked by authors. In this paper, we adopt the convention by which negative ΔΔG values indicate destabilizing variations. Interestingly, the dataset has been built to be non-redundant at 25% sequence identity with respect to datasets routinely used for training tools available in literature, including the S2648 (Dehouck *et al*., 2011) and the VariBench dataset (Nair and Vihinen, 2013). This enables a fair comparison with most state-of-the-art tools. Variations included in S669 are provided in relation to PDB chains. In this work, since DDGemb adopts protein language models for input encoding, we mapped all variations on full-length UniProt (https://www.uniprot.org/) sequences using SIFTS (Dana *et al*., 2019).

#### 2.1.2. Training set: the S2450 dataset

To build our training set, we started from the well-known and widely adopted S2648 dataset (Dehouck *et al*., 2011), containing 2648 single-point variations on 131 different proteins. Associated experimental ΔΔG values are retrieved from the ProTherm database (Bava, 2004) and were manually checked and corrected to avoid inconsistencies. Differently from previous works adopting the same dataset (Dehouck *et al*., 2011; Fariselli *et al*., 2015), in which variations are directly mapped on PDB chain sequences, in this work, we adopted full-length protein sequences from UniProt. To this aim, we used SIFTS (Dana *et al*., 2019) to map PDB chains and variant positions on corresponding UniProt sequences.

Homology reduction of the S669 dataset against S2648 was originally performed in (Pancotti *et al*., 2022) considering only PDB-covered portions of the sequences. This procedure does not guarantee to detect all sequence similarity on full-length sequences. For this reason, in this work, we compared UniProt sequences in S669 and S2648, removing from the training set those having more than 25% sequence identity with any sequence in the test set (S669). Overall, 18 sequences were removed from S2648, accounting for 198 single-point variations. This reduced dataset is then referred to as S2450 throughout the entire paper.

The S2450 dataset was adopted here to perform 5-fold cross-validation. To this aim, we implemented the stringent data split procedure described in (Fariselli *et al*., 2015), by which all variations occurring on the same protein are put in the same cross-validation subset and proteins are divided among subsets taking into consideration pairwise sequence identity (setting a threshold to 25%). In this way, during cross-validation, no redundancy is present between protein sequences included in training and validation sets.

By construction, the S2450 dataset is unbalanced toward destabilizing variations (i.e., negative ΔΔG values). To balance the dataset, to reduce the bias towards destabilizing ΔΔG values, and to improve the model capability of predicting stabilizing variations, we exploited thermodynamic reversibility of variations, by which ΔΔG(A -> B) = -ΔΔG(B -> A) (Capriotti *et al*., 2008). Using the reversibility property, the set of variations can be artificially doubled to include reverse variations, switching the sign of experimental ΔΔG values.

#### 2.1.3. Multiple variations: the reduced PTmul dataset

We also adopted a dataset for testing the DDGemb on the prediction of ΔΔG upon multi-point variations. The dataset, referred to as PTmul, has been introduced in (Montanucci *et al*., 2019): it comprises 914 multi-point variations on 91 proteins. However, the original PTmul dataset share a high level of sequence similarity when compared to our S2450 training dataset. In order to perform a fair evaluation of the performance, we excluded from PTmul all proteins that are similar to any protein in our S2450 training set. After this reduction step, we retained 82 multi-point variants occurring in 14 proteins. Although the number of variants is significantly reduced if compared to the original dataset, using this homology reduction procedure ensures a fair evaluation of the different methods. The reduced dataset is referred to as PTmul-NR.

### 2.2. The DDGemb method

An overview of the DDGemb deep-learning model is shown in Figure 1. The architecture comprises two components: i) the input encoding and ii) the ΔΔG prediction model. In the next section we describe in detail both components.

**Figure 1.**
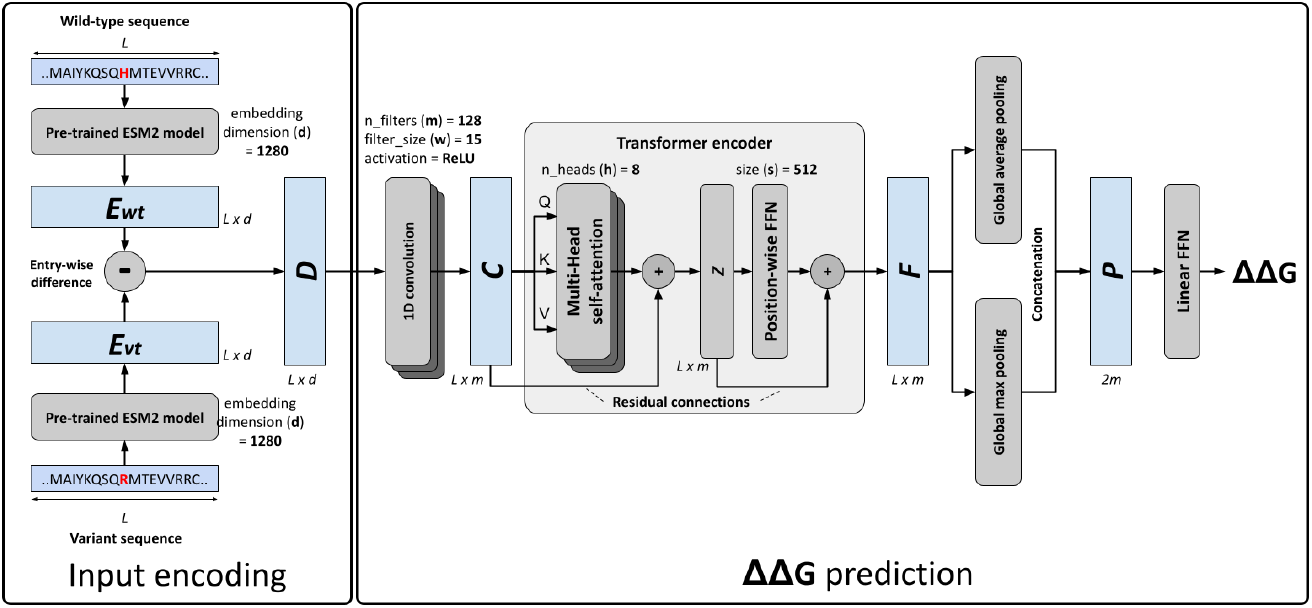
The DDGemb model architecture.

#### 2.2.1. Input encoding

For encoding a single residue variation, we start from the wild-type and the variant protein sequences. The latter is derived from the former upon either single-point or multi-point variations. In the first step, the two sequences, both of length *L*, are encoded using the ESM2 protein language model (pLM) (Lin *et al*., 2023; Rives *et al*., 2021). Among the different models available and after input encoding optimization (see Results), here we adopted the medium-size 33-layers model with 650M parameters and trained on the UniRef50 database. This model provides residue-level embeddings of dimension 1280 and represents a good trade-off between representation expressivity and computational requirements. For generating embeddings, we adopted the ESM2 package available at https://github.com/facebookresearch/esm.

The application of the ESM2 pLM provides two *L* × *d* matrices, named *E*_*wt*_ and *E*_*vt*_, representing the residue-level embeddings of the wild-type and variant sequences (derived either from a single- or multi-point variation), respectively. A single *L* × *d* matrix *D* encoding the variation is then generated computing the element-wise difference of *E*_*wt*_ and *E*_*vt*_:

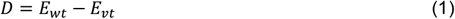

The matrix *D* is used as input for the downstream ΔΔG prediction architecture.

#### 2.2.2. The Transformer based ΔΔG prediction network

The remaining part of the DDGemb architecture is devised to predict a ΔΔG value starting from the input matrix *D* encoding the protein variant. The hyperparameters of the final model were optimized in cross-validation, according to the different configurations reported in Table 1 (see Results section). After optimization, the final selected model is Model4 (Table 1), described in the following.

**Table 1.**
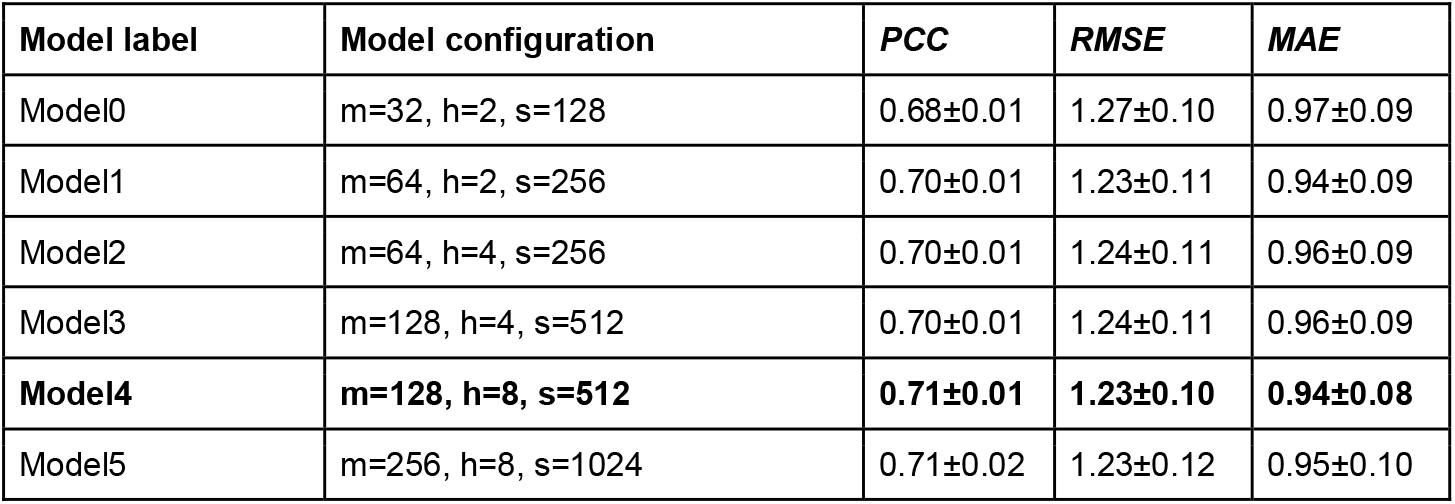
Five-fold cross-validation results of different ΔΔG prediction architectures

The input matrix *D* is firstly processed by a one-dimensional (1D) convolution layer comprising *m*=128 filters of size *w*=15, with ReLU activation functions. The 1D-convolution layer provides a way of projecting the higher-dimensional input data into a lower-dimensional space of size *m*, extracting local contextual information through a series of sliding filters of width *w*. The output of the 1D-convolution is a matrix C of dimension *L* × *m*.

The matrix *C* is then passed through a Transformer encoder layer (Vaswani *et al*., 2017), consisting of a cascading architecture including a multi-head attention layer with 8 attention heads (*h*), residual connections, and a position-wise feedforward network (FFN). The Transformer encoder is responsible for computing self-attention across the input sequence, producing in output a representation of the input taking into consideration the relations among the different positions of the input sequence.

The architecture adopted here is directly derived from the original Transformer definition (Vaswani *et al*., 2017). Formally, given the input sequence *C* of dimension *L* × *m*, each head *i* of the multi-head attention layer adopts three matrices of learnable weights, called 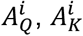 and 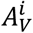 each having dimension *m* × *r*, where *r* = *m/h* (*r* is equal to 16 in our case) and *h* is the number of attention heads (here set to 8). The input matrix *C* is firstly projected using the 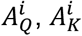 and 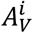 as follows:

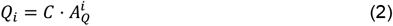

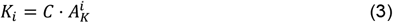

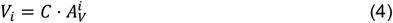

where ⋅ denotes the matrix product operator.

Then, for each head *i*, an attention output *Z*_*i*_ of dimension *L* × *r* is computed as follows:

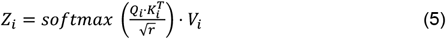

The different *Z*_*i*_ from the different attention heads are then concatenated, multiplied with an output weight matrix *A*_*O*_ of dimension *m* × *m*, and the result added to the input *C* matrix by residual connection:

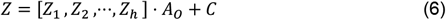

where [ ] denotes the concatenation operator (by rows) and the Z output matrix has dimension *L* × *m* (as the input matrix *C*).

The final Transformer encoder output *F* of dimension *L* × *m* is computed by independently applying a position-wise Feed-Forward Network (FFN) to each position 1 ≤ *j* ≤ *L* of *Z*, and residual connection addition. In other words, each row *f*_*j*_ of F is computed as follows:

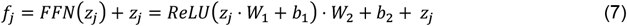

where ReLU is the activation function defined as *g*(*x*) = *max*(0, *x*) and *W*_1_, *b*_1_, *W*_2_, *b*_2_, are position-independent weight parameters and biases of FFN, having, respectively, dimensions *m* × *s*, 1 × *s, s* × *m* and 1 × *m*. Here, we set the dimensionality *s* of the hidden layer of the position-wise FFN to 512.

The output matrix *F* of the Transformer encoder is collapsed to two vectors of dimension *m* by means of Global Average and Max Pooling layers, denoted as *p*_*ave*_ and *p*_*max*_, respectively, acting on the first dimension *L* of the matrix *F*:

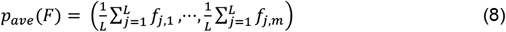

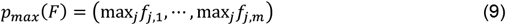

The pooled vectors are then concatenated into a single vector *P* of size 2*m*:

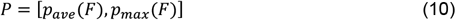

where [⋅] denotes the concatenation operator.

*P* is finally processed by a linear FFN parametrized by a weight vector *w*_*o*_ and bias *b*_*o*_, producing in output the predicted ΔΔG value *ŷ*:

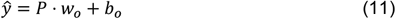

#### 2.2.3. Model training and implementation

Given a dataset of *N* protein variants *D* = {*D*^1^, *D*^2^,⋯, *D*^*N*^}, corresponding target ΔΔG values *Y* = {*y*^1^, *y*^2^,⋯, *y*^*N*^}, and model predictions *Ŷ* = {*ŷ*^1^, *ŷ*^2^,⋯, *ŷ*^*N*^}, training is carried out minimizing the Mean Squared Error function on training data:

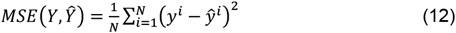

The optimization is carried out with gradient descent and adopting the Adam optimizer (Kingma and Ba, 2017). The training data are split into mini batches of size 128. Training is performed for 500 epochs and stopped when error starts decreasing on a subset of the training set used as validation data (early stopping).

The training procedure and the model itself were implemented using the PyTorch Python package (https://pytorch.org). All experiments were carried out on a single machine equipped with two AMD EPYC 7413 CPUs with 48/96 CPU cores/threads and 768GB RAM.

### 2.3. Scoring performance

To score the performance of the different approaches, we use the following well-established scoring indexes. In what follows, *e* and *p* are experimental and predicted ΔΔG values, respectively, while *p*^*dir*^ and *p*^*rev*^ are predicted ΔΔG for direct and corresponding reverse variations, respectively.

The Pearson’s correlation coefficient (PCC) between *e* and *p* is defined as:

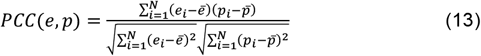

where *ē* and 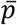 are average experimental and predicted ΔΔG values, respectively.

The Root Mean Square Error (RMSE) between *e* and *p* is defined as:

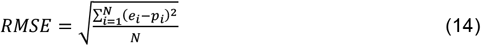

The Mean Absolute Error (MAE) between *e* and *p* is defined as:

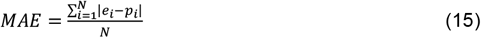

To score anti-symmetry properties of the different tools we adopted two additional measures defined in literature (Pucci *et al*., 2018).

The Pearson’s correlation between *p*^*dir*^ and *p*^*rev*^, referred to as *PCC*_*d*−*r*_, defined as:

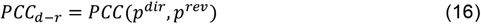

The anti-symmetry bias *⟨δ⟩* defined as:

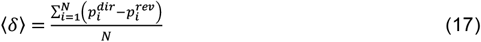

## 3. Results

### 3.1. Cross-validation results on the S2450 dataset

In a first experiment, we performed 5-fold cross-validation on the S2450 dataset. To this aim, we adopted the most stringent data split procedure proposed in (Fariselli *et al*., 2015), which consists in retaining all variations occurring in the same protein within the same cross-validation subset and in confining proteins with more than 25% sequence identity in the same subset. Sequence comparison was performed using full-length UniProt sequences.

We performed a cross-validation to optimize the hyperparameters defining the model architecture. Specifically, we trained six models with different architectures as reported in Table 1, varying three hyperparameters: the number of filters in the 1D-convolutional layer (*m*), the number of Transformer encoder attention heads (*h*) and the dimension of hidden layer in the Transformer encoder module (*s*). The best model architecture was chosen according to the average cross-validation *PCC* between experimental and predicted ΔΔG. Results are presented in Table 1.

Considering both average *PCC, RMSE* and *MAE* values and the corresponding standard deviations, the highest performance is obtained using the architecture Model4, including 128 1D-convolutional filters, 8 Transformer encoder attention heads and 512 hidden units in the Transformed encoder FFN output. This configuration has been then chosen as the final model.

### 3.2. Prediction of single-point variations on the S669 dataset

We compared DDGemb with several state-of-the-art methods introduced in the past years using the common benchmark dataset S669. Results for 21 different methods were taken from (Pancotti *et al*., 2022), except DDGemb, presented in this work, PROSTATA (Umerenkov *et al*., 2023) and THPLM (Gong *et al*., 2023), whose results were extracted from the respective papers. Scored methods include eight sequence-based predictors, namely INPS (Fariselli *et al*., 2015), ACDC-NN-Seq (Benevenuta *et al*., 2021), DDGun (Montanucci *et al*., 2019), I-Mutant3-Seq (Capriotti *et al*., 2005), SAAFEC-SEQ (Li *et al*., 2021), MUPro (Cheng *et al*., 2006), PROSTATA (Umerenkov *et al*., 2023), and THPLM (Gong *et al*., 2023), and fifteen structure-based methods, ACDC-NN (Benevenuta *et al*., 2021), PremPS (Chen *et al*., 2020), DDGun3D (Montanucci *et al*., 2019), INPS-3D (Savojardo *et al*., 2016), ThermoNet (Li *et al*., 2020), MAESTRO (Laimer *et al*., 2015), Dynamut (Rodrigues *et al*., 2018), PoPMuSiC (Dehouck *et al*., 2011), DUET (Pires *et al*., 2014a), SDM (Worth *et al*., 2011), mCSM (Pires *et al*., 2014b), Dynamut2 (Rodrigues *et al*., 2021), I-Mutant3-3D (Capriotti *et al*., 2005), Rosetta (Kellogg *et al*., 2011) and FoldX (Schymkowitz *et al*., 2005). Results are listed in Table 2.

**Table 2.**
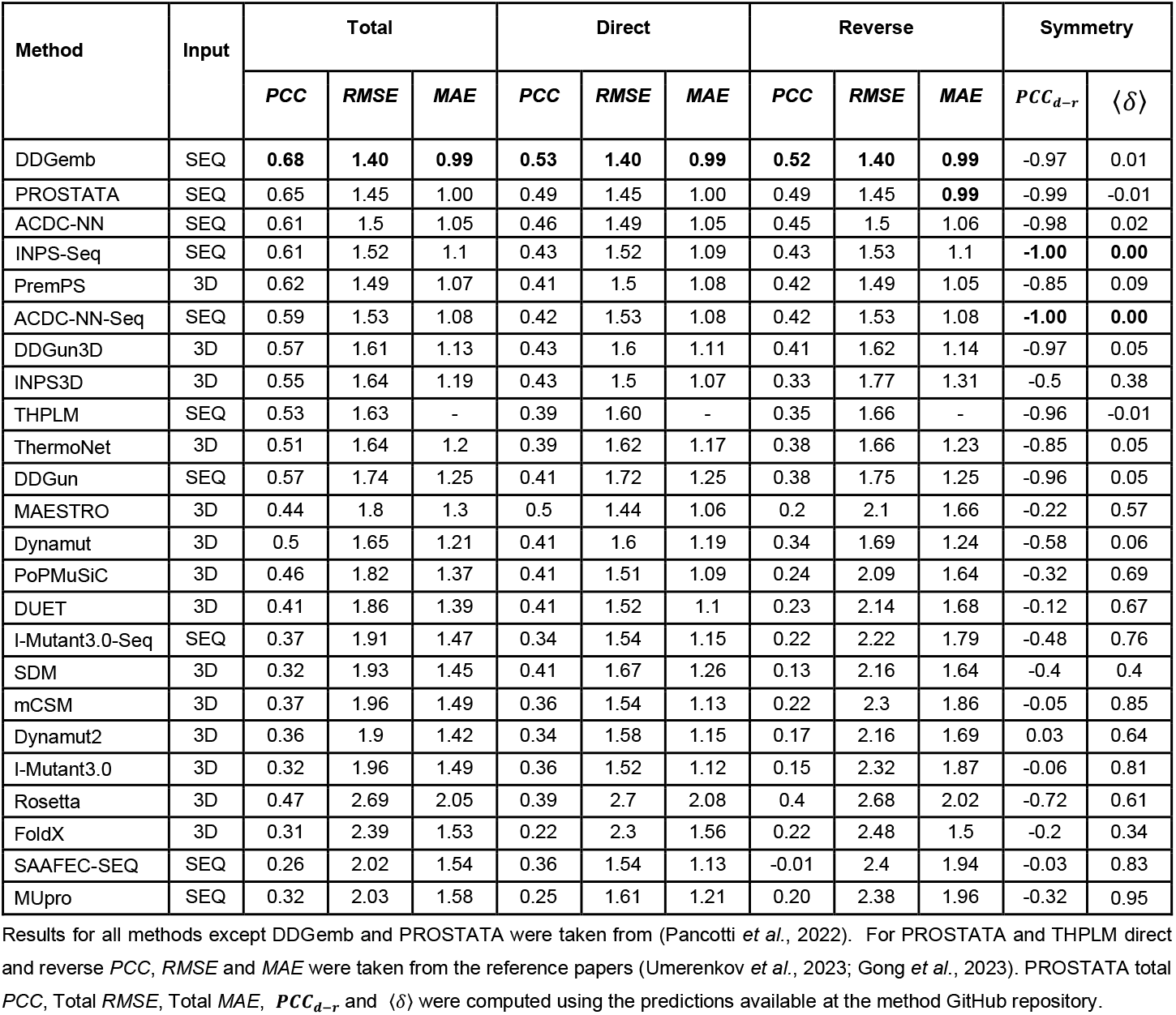
Comparative benchmark of different sequence- and structure-based methods on the S669 independent test set

For each method we report *PCC, RMSE* and *MAE* computed considering i) all variations (both direct and reverse) in the dataset (columns under “Total”), ii) only direct variations (columns under “Direct”) and iii) only reverse variations (columns under “Reverse”). In addition, we computed *PCC*_*d*−*r*_ and *⟨δ⟩*.

On the S669 dataset, DDGemb reports the highest *PCC, RMSE and MAE* values (Total, Direct and Reverse). Our DDGemb overall scores as the top-performing tool in this benchmark, significantly overpassing both structure- and sequence-based methods.

### 3.3. Prediction of multi-point variations

We finally tested DDGemb in the prediction of multi-point variations using the PTmul-NR dataset. This allowed us to directly compare with other methods such as DDGun/DDGun3D (Montanucci *et al*., 2019), MAESTRO (Laimer *et al*., 2015) and FoldX (Schymkowitz *et al*., 2005). Results are listed in Table 3.

**Table 3.**
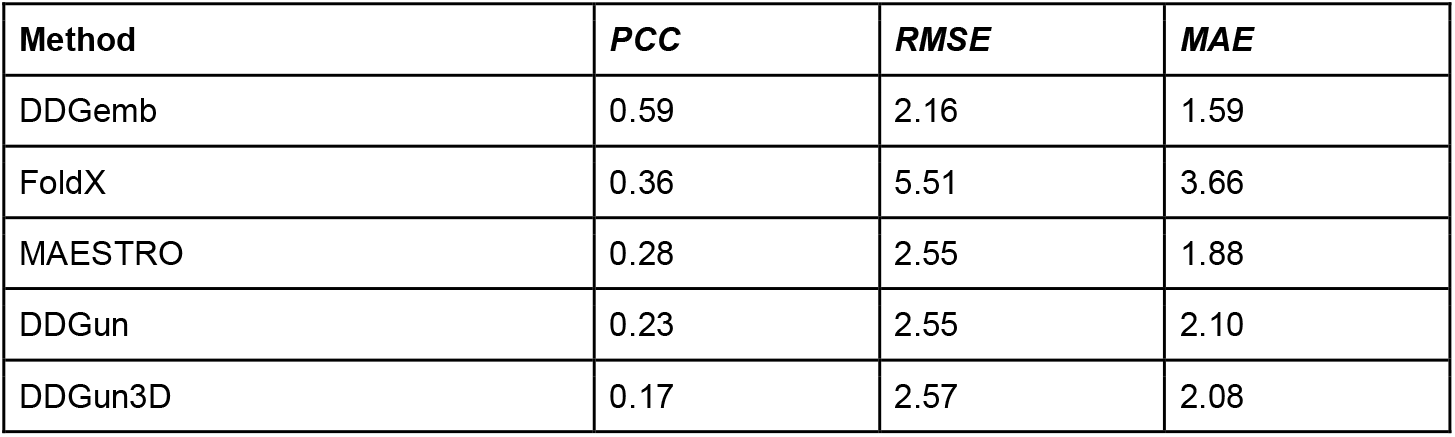
Comparative benchmark of different methods on multi-point variations from the PTmul-NR dataset

On the PTmul-NR dataset, DDGemb significantly outperforms DDGun, DDGun3D, FoldX and MAESTRO on the prediction of multi-point variations, achieving the highest correlation coefficient of 0.59, and the lowest RMSE and MAE values of 2.16 and 1.59, respectively.

These results suggest that DDGemb can be effectively used for assessing with high accuracy the impact of multi-point variations on protein stability. Remarkably, our method has been trained using only single-point variants, suggesting the ability of the proposed approach to generalize on multi-point variants as well.

## 4. Conclusions

In this work we present DDGemb, a novel method based on protein language models and Transformers to predict protein stability change (ΔΔG) upon single- and multi-point variations. Our method has been trained on a high-quality dataset derived from literature and tested using recently introduced benchmark datasets of thermodynamic data for single- and multi-point variations. In all the benchmarks, DDGemb reports performances that are superior to the state of art, outperforming both sequence- and structure-based methods and achieving an overall *PCC* of 0.68 on single-point variations. Moreover, on multi-point variations, our method reports a *PCC* of 0.59, which is significantly higher than the one achieved by the second top-performing approach, FoldX, reporting PCC equal to 0.36. Our study suggests the relevance of a transformer architecture specifically fine-tuned to predict the ΔΔG upon variation, in combination with numerical representations provided by protein language models.

